# Paragraph - Antibody Paratope prediction using Graph Neural Networks with minimal feature vectors

**DOI:** 10.1101/2022.06.10.495640

**Authors:** Lewis Chinery, Newton Wahome, Iain Moal, Charlotte M. Deane

## Abstract

1

**Summary:** The development of new vaccines and antibody therapeutics typically takes several years and requires over $1bn in investment. Accurate knowledge of the paratope (antibody binding site) can speed up and reduce the cost of this process by improving our understanding of antibody-antigen binding. We present Paragraph, a structure-based paratope prediction tool that outperforms current state-of-the-art tools using simpler feature vectors and no antigen information.

**Availability:** Source code is freely available at www.github.com/oxpig

**Supplementary information:** Supplementary data are available at *bioRxiv* online.

## 2 Introduction

Antibodies can bind to and neutralise their target antigens with high specificity and affinity. Their high specificity and low immunogenicity means antibodies have grown to dominate the therapeutic marketplace[1]. Current experimental methods to determine how an antibody and antigen bind, and hence the potential neutralising effect, are slow and costly[2]. Computational docking tools[3, 4, 5] have been developed to predict binding, offering a cheaper, faster alternative. However, current computational tools often fail to accurately recapitulate antibody-antigen binding[6] and are generally not feasible for use in a truly high throughput fashion[7]. Predicting the binding site of an antibody can both allow better estimation of its bound pose[8] and indicate key residues to mutate to change binding properties[9].

Parapred[10] is a widely-used and freely available sequence-based paratope prediction tool that uses convolutional and recurrent neural networks. More recent attempts at paratope prediction e.g. PECAN[11], have brought in structural information, representing the antibody structure as a graph. PECAN requires both antibody and antigen structures as input into a graph convolution attention network, but based on that input outperforms Parapred (PR AUCs of 0.675 and 0.646 respectively, Table 1).

**Table 1:**
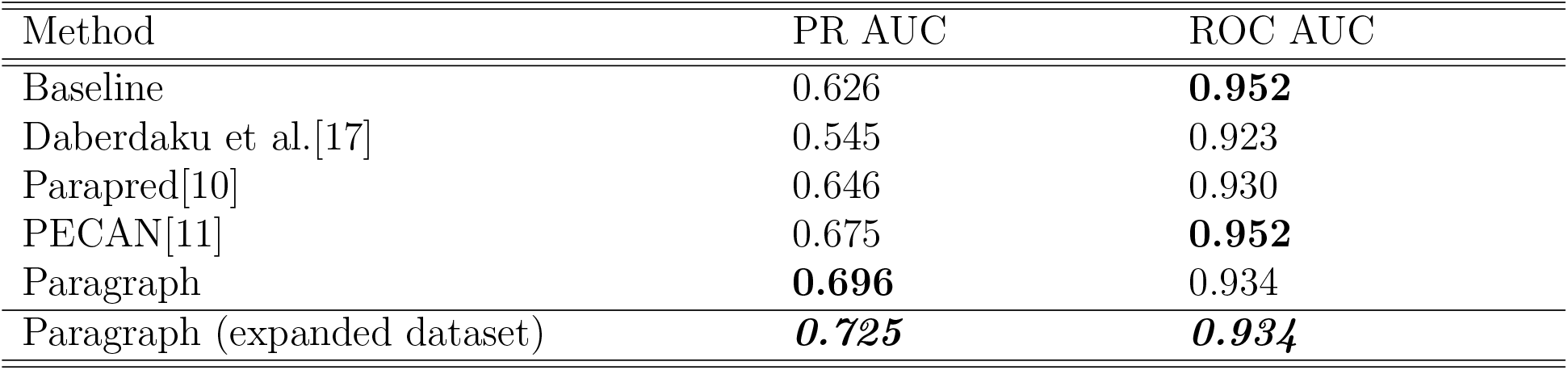
Comparison of paratope prediction methods. All are evaluated on model structures over the *F*_*v*_ region. All results use PECAN’s dataset unless stated otherwise. Daberdaku et al.’s dataset includes 11 additional complexes with non-protein antigens. Daberdaku et al. and PECAN’s performance have been taken from the original papers (PECAN’s numbers are approximated from the figures provided). We have retrained Parapred using PECAN’s dataset and extended the results over the *F*_*v*_ by predicting zero for all additional residues, similar to our method. Our baseline (see Results section) shows the importance of considering PR AUC in this class imbalanced problem.

In this paper we take advantage of methods that are able to accurately and rapidly predict antibody structures[12] to build a structure-based paratope predictor, Paragraph. As crystal structures will only be available for a tiny number of the antibodies of interest[13, 14], it is essential for structure-based paratope predictors to work on models. Paragraph takes as input the modelled structure of an antibody and, using equivariant graph neural network layers[15], rapidly predicts the probability of residues to belong to the paratope. Paragraph outperforms the current state-of-the-art paratope predictor, PECAN, achieving a PR AUC of 0.696 when trained on the same dataset.

## 3 Data

Paragraph was trained and tested on the same complexes used by PECAN[11]. This dataset contains 460 antibody-antigen complexes - 205 complexes are used for training, 103 for validation, and 152 for testing. The dataset includes only complexes with paired heavy and light chains, sub 3Å resolution, and protein antigens.

To take advantage of the ever increasing amount of structural data now available, we also trained Paragraph on a larger dataset. This new dataset was extracted from the Structural Antibody Database (SAbDab)[13] on 31/03/2022 and includes 1,086 complexes which we divide into train, validation, and test sets using a 60-20-20 split. Full details of both datasets are given in the SI. To build our final models, we use ABodyBuilder[16] to model the framework region and ABlooper[12] to model the CDR loops, not allowing the use of sequence identical templates. Previous methods have chosen to search either the entire *F*_*v*_ region for paratope residues[11, 17] or the CDR±2 only[10]. To allow direct comparison to all methods, Paragraph has been trained on the smaller search area (using IMGT-defined[18] CDR loops), with the predictions then extended to the entire *F*_*v*_ by predicting non-binding for all residues outside the CDR±2. In our Results we present the *F*_*v*_ region performance; CDR±2 results are given in the SI. All PDB codes and heavy, light, and antigen chain IDs used in both datasets can be found at www.github.com/oxpig.

## 4 Methods

To maximise the performance of Paragraph on model structures, we trained on a combination of both crystal and model structures and validated on model structures only.

For both crystal and model structures, we generate a graph-representation of the antibodies by describing each residue in the CDR±2 as a single node. Directionless edges are defined to exist between nodes separated by 10Å or less. Each node has a 22D feature vector comprised of a 20D onehot encoding of the amino acid type and a 2D onehot encoding of the chain type. Following previous methods[10, 11], residues were labelled as binding if any heavy atom was within 4.5Å of an antigen heavy atom.

To predict the binding of each residue, a series of six graph and three linear layers were used. The graph layers were adapted from LucidRain’s[19] implementation of the architecture described in E(n) Equivariant Graph Neural Networks[15]. In our network the coordinates of our nodes are fixed. Further details of our architecture and training can be found in the SI.

To extend our predictions over the entire *F*_*v*_ we predict zero for all residues outside the CDR±2. This simple extension works well as only 1% of residues outside the CDR±2 are classed as binding in both datasets. We used this approach instead of training over and predicting the entire *F*_*v*_ region as the large class imbalance (10:1) resulted in high instability.

## 5 Results

The top row of Table 1 gives the performance of a simple baseline which measures the proportion of residues found to bind the antigen in our training set at each sequence position. We then predict this same proportion for every corresponding position in our test set. This baseline achieves the highest ROC AUC of all methods, demonstrating the importance of considering the area under the precision-recall curve, PR AUC, for this class imbalanced problem.

Paragraph outperforms both PECAN[11] and Parapred[10], the best performing freely available methods (Table 1). Paragraph is also faster than existing methods, taking approximately 0.1s to predict the paratope when fed a PDB file, 50 times faster than Parapred. Finally, Paragraph also requires no antigen information, meaning it can be deployed in wider use cases than PECAN. The performance of Paragraph is further boosted by the use of the larger dataset of structures now available (Table 1).

Additional details and breakdowns of Paragraph’s performance can be found in the SI, including different metrics, search areas, and datasets.

## 6 Discussion

Due to the large class imbalance present in the task of paratope prediction, care must be taken when selecting evaluation metrics as some metrics may give misleadingly high performance. We show that our simple baseline achieves the highest ROC AUC (0.952) of all paratope prediction methods tested (Table 1), highlighting the need to use PR AUC when evaluating this class imbalanced problem.

We also stress the importance of using model, not crystal, structures when evaluating performance due to the lack of readily available crystal structures in research settings. Were crystal structures to be used for testing in place of model structures, Paragraph’s PR AUC on our expanded dataset increases from 0.725 to 0.757 (SI).

Using equivariant graph neural networks, simple feature vectors, and no antigen information, we achieve paratope prediction performance above the current state-of-the-art, PECAN, which requires structures of both the antibody and antigen. On the PECAN dataset, Paragraph achieves a PR AUC of 0.696 and the performance increases further when you use an expanded dataset, suggesting as more data becomes available, further improvements may be achieved.

In addition, when considering only ABlooper’s 50% most confident models, Paragraph achieves a PR AUC of 0.763 on our expanded dataset (SI). Further advancements in antibody structure prediction should therefore also result in more accurate paratope prediction. The correlation of Paragraph’s performance with ABlooper’s model confidence also enables users to better understand when their predicted paratope is likely to be accurate.

## Supporting information

Paragraph Supplementary Information

## Funding

This work was supported by the Biotechnology and Biological Sciences Research Council (BB-SRC) and GlaxoSmithKline (GSK).

## Conflict of Interest

none declared.

## Notes

https://github.com/oxpig

### Competing Interest Statement

The authors have declared no competing interest.

