## Supplementary material for "Paragraph - Antibody Paratope prediction using Graph Neural Networks with minimal feature vectors": Paragraph Supplementary Information

#### 1 Datasets

##### 1.1 PECAN dataset

PECAN's[1] dataset contains 460 antibody-antigen complexes - 205 complexes are used for training, 103 for validation, and 152 for testing. PECAN's dataset is a subset of the 472-complex dataset compiled by Daberdaku et al.[2]. The reduced 460 complex dataset contains only protein and peptide antigens. CSVs containing the PDB codes and heavy, light and antigen chain IDs used in our training, validation and testing can be found at www.github.com/oxpig.

In our research, we identified issues with certain structures in the PECAN dataset - a subset of these are shown below. These issues, along with the large growth in structural data over the past three years, inspired us to create our Expanded dataset (Section 1.2).

##### 1.1.1 1IGC - antigen not binding $F_v$

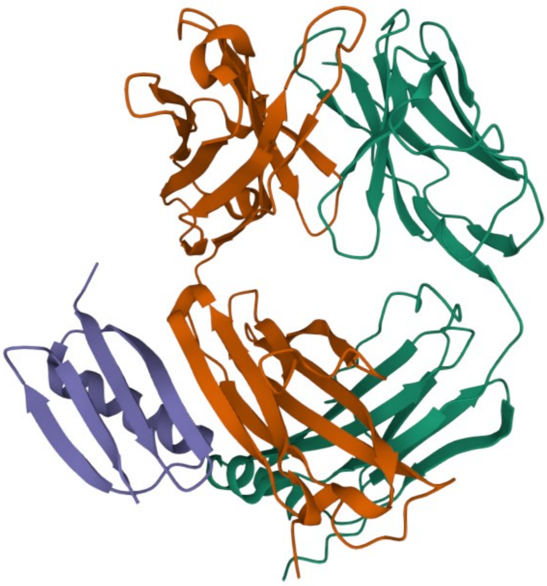

Figure S1: A cartoon representation of 1IGC taken from https://www.rcsb.org/. The heavy chain is shown in orange, light chain in green, and antigen in purple. The antigen binds entirely outside of the $F_v$ region.

PDB structure 1IGC is found in the PECAN train set and includes an antigen bound to the constant region of the $F_{ab}$. No contacts exist in the $F_v$ region (Figure S1).

##### 1.1.2 4ERS - antigen misplaced in PDB file

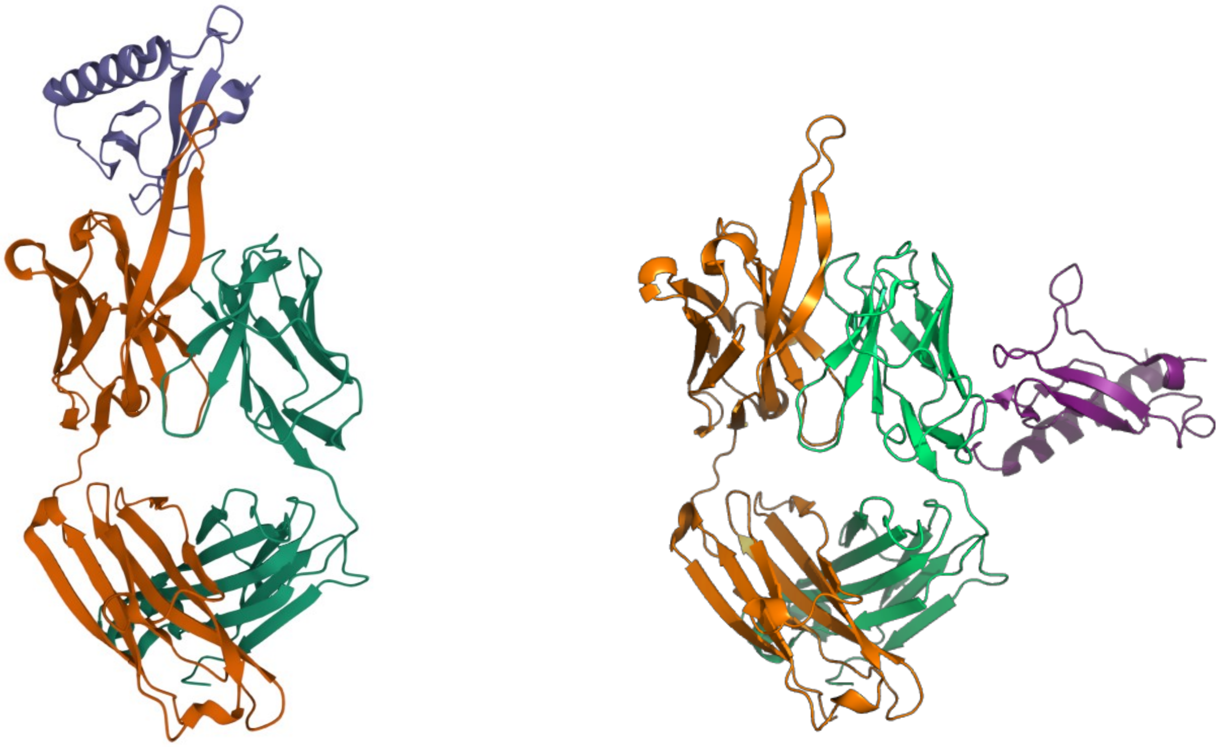

Figure S2: Left image shows a cartoon representation of 4ERS taken from https://www.rcsb.org/. The heavy chain is shown in orange, light chain in green, and antigen in purple. The right image was created in PyMOL where colours have been chosen to approximately match the RCSB image. The antigen clearly appears in two different positions in the two figures.

PDB structure 4ERS is found in the PECAN validation set. On the RCSB webpage the true binding confirmation of antibody and antigen can be found. However, in the PDB file the incorrect antigen has been paired with the antibody (Figure S2).

##### 1.1.3 1TZI - One chain of antigen homodimer removed in PDB file

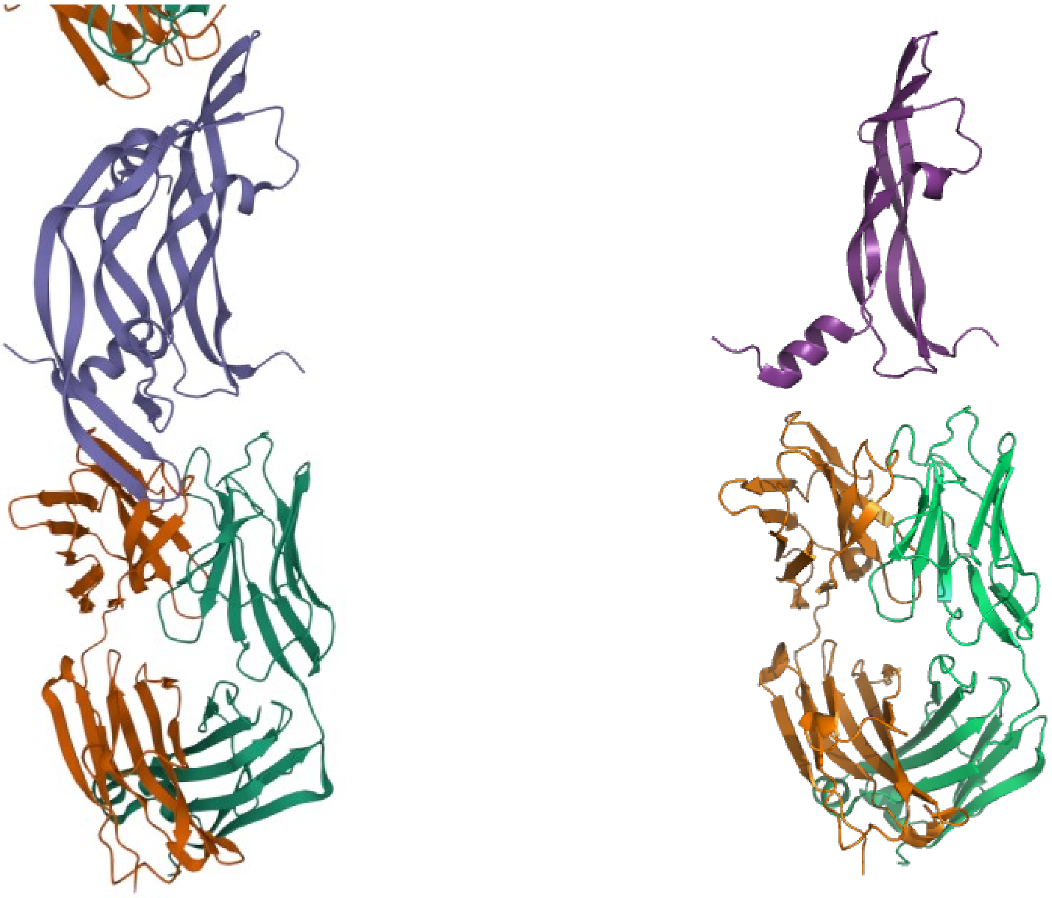

Figure S3: Left image shows a cartoon representation of 1TZI taken from https://www.rcsb.org/. The heavy chain is shown in orange, light chain in green, and antigen in purple. The right image was created in PyMOL where colours have been chosen to approximately match the RCSB image. The antigen chain that dominates binding in the left figure is not present in the PDB file.

PDB structure 1TZI is found in the PECAN validation set. On the RCSB webpage it is clear that the antibody is binding both chains of the homodimer antigen. However, in the PDB file the dominantly binding chain of the homodimer has been removed in this file only. Four contacts exist to the $F_v$ region (Figure S3).

#### 1.2 Expanded dataset

To take advantage of the ever increasing amount of structural data now available, we also trained Paragraph on a larger dataset. This new dataset was extracted from the Structural Antibody Database (SAbDab)[3] on 31/03/2022 and includes 1,086 complexes which we divide into train, validation, and test sets using a 60-20-20 split. To generate this new dataset we included only paired heavy and light chain X-ray complexes with 3Å resolution or below. We allow both protein and peptide antigens, as defined by SAbDab.

To overcome data quality issues observed in the PECAN dataset we require that the $F_v$ contains 10 or more binding residues and that 50% or more of these binding residues lie in the CDR loops plus two residues on either side (CDR+2). These two requirements remove just 5% of the dataset.

Finally, using CD-HIT[4], we ensure no two antibodies share over 95% heavy and light chain sequence identity and we remove any structures ABodyBuilder[5] cannot model using a 95% identity threshold (where no suitable templates were found for homology modelling of the framework region).

The Expanded training, validation and test set data can be found at www.github.com/oxpig.

### 2 Paragraph architecture and training

To maximise the performance of Paragraph on model structures, we trained on a combination of both crystal and model structures and validated on model structures only.

For both crystal and model structures, we generate a graph-representation of the antibodies by describing each residue in the CDR±2 as a single node. The nodes are defined to have coordinates given by the $C_\alpha$ atoms. Directionless edges are defined to exist between nodes separated by 10Å or less. Each node has a 22D feature vector comprised of a 20D onehot encoding of the amino acid type and a 2D onehot encoding of the chain type. Following previous methods[6, 1], residues were labelled as binding if any heavy atom was within 4.5Å of an antigen heavy atom.

To predict the binding of each residue, a series of six graph and three linear layers were used. The graph layers were adapted from LucidRain's[7] implementation of the architecture described in E(n) Equivariant Graph Neural Networks[8]. In our network the coordinates of our nodes are fixed.

All graph layers have input and output feature dimensions of 22 and utilise skip connections. The linear layers have output dimensions of 10, 10, and 1. Following each graph layer the non-linear Hardtanh activation function is applied. Hardtanh activation is also applied following the first two linear layers, while a Sigmoid function is applied to the output of the final layer to yield a probability between zero and one.

Paragraph was trained for 16 different random seeds, each for 300 epochs. Batch sizes of one were used and gradients were accumulated across 16 batches. The Binary Cross Entropy (BCE) loss was calculated between the residue labels and predictions. This loss was then optimised using Adam stochastic gradient descent with a learning rate of 0.001. An imbalance weighting of three was used to account for the class imbalance between residues classed as binding the antigen (27%) and those classed as not binding (73%). For each seed, weights were saved that delivered the lowest BCE loss on the validation dataset. The weights that resulted in the largest area under the precision-recall curve (PR AUC) for the validation set were then used to predict the paratope for models of our test set.

To extend our predictions over the entire $F_v$ we predict zero for all residues outside the CDR±2. This simple extension works well as only 1% of residues outside the CDR±2 are classed as binding in both datasets. We used this approach instead of training over and predicting the entire $F_v$ region as the large class imbalance (10:1) resulted in high instability.

### 3 Paragraph performance on both datasets

PR AUC is the preferred metric for evaluating highly class imbalanced problems. Below we also give Paragraph's performance evaluated using other popular metrics to allow quick comparison to other methods (Table S1). Results are provided on ABodyBuilder+ABlooper[5, 9] model structures of our test sets and evaluated over the entire $F_v$ region. In agreement with Parapred[6], thresholds for calculating the F-score and Matthews correlation coefficient (MCC) are obtained by maximising Youden's index[10].

| Dataset | PR AUC | ROC AUC | F-score | MCC |
|---|---|---|---|---|
| PECAN | 0.696 | 0.934 | 0.685 | 0.654 |
| Expanded | 0.725 | 0.934 | 0.696 | 0.669 |

Table S1: Performance of Paragraph across the entire $F_v$ on model structures.

### 4 Paragraph performance over the CDR loops plus-minus 2 extra residues

Previous paratope prediction methods such as Parapred[6] have searched for paratope residues in only the CDR loops plus two residues on either side. This restriction reduces the search area by a factor of three compared to the entire $F_v$ region while still retaining ~92% of all paratope residues. (The remaining 8% of paratope residues make up just 1% of all residues outside this search area.) The smaller class imbalance faced in this problem leads to improved training performance evaluated using the PR AUC. In contrast to Parapred, which uses Chothia defined CDR loops, we use the IMGT[11] definition. These two definitions share 98% of residues. All results are provided on ABodyBuilder+ABlooper[5, 9] model structures of our test sets (Table S2). In agreement with Parapred[6], thresholds for calculating the F-score and Matthews correlation coefficient (MCC) are obtained by maximising Youden's index[10].

| Dataset | PR AUC | ROC AUC | F-score | MCC |
|---|---|---|---|---|
| PECAN | 0.730 | 0.891 | 0.705 | 0.584 |
| Expanded | 0.765 | 0.907 | 0.717 | 0.606 |

Table S2: Performance of Paragraph across the CDR±2 on model structures.

### 5 Paragraph performance on crystal and model structures

Until recently, structure-based paratope prediction methods were limited in their usefulness by the lack of structural data or accurate models. However, recent advancements in antibody modelling[5, 9] mean structure-based paratope prediction methods can now be used widely.

The below table (Table S3) shows that if we select only ABlooper's most confident models (those with the lowest H3 decoy diversity), we can match the performance observed when evaluating Paragraph on crystal structures. All results are provided for our Expanded test set and are evaluated over the entire $F_v$ region. In agreement with Parapred[6], thresholds for calculating the F-score and Matthews correlation coefficient (MCC) are obtained by maximising Youden's index[10].

| Trained on | PR AUC | ROC AUC | F-score | MCC |
|---|---|---|---|---|
| Crystals | 0.757 | 0.937 | 0.719 | 0.692 |
| Models (all) | 0.725 | 0.934 | 0.696 | 0.669 |
| Models (top 75%) | 0.742 | 0.937 | 0.713 | 0.687 |
| Models (top 50%) | 0.763 | 0.939 | 0.727 | 0.700 |
| Models (top 25%) | 0.800 | 0.947 | 0.755 | 0.731 |

Table S3: Performance of Paragraph across the entire $F_v$ on both crystal and model structures using our Expanded dataset.

### 6 Paragraph performance on individual heavy and light chains

When trained on paired data, Paragraph learns information pertinent to both the heavy and light chains. This means that the performance falls when predicting paratope residues for antibodies where data is only available for one chain. To overcome this, Paragraph has also been trained on only heavy and only light chains. These single chain weights are made available with the source code at www.github.com/oxpig. Paragraph's performance when trained and tested on only the heavy or light chains of our Expanded dataset are provided below (Table S4). In agreement with Parapred[6], thresholds for calculating the F-score and Matthews correlation coefficient (MCC) are obtained by maximising Youden's index[10].

The performance of Paragraph's light chain predictions are worse than the heavy chain predictions due to the greater class imbalance - fewer light chain residues belong to the paratope overall, and there are a few examples in the dataset where no residues at all belong to the positive class.

As expected, neither individual chain's performance exceeds the paired performance of Paragraph when evaluated on the chain in question.

| Chain | PR AUC | ROC AUC | F-score | MCC |
|---|---|---|---|---|
| Heavy | 0.737 | 0.943 | 0.713 | 0.681 |
| Light | 0.637 | 0.910 | 0.630 | 0.608 |

Table S4: Performance of Paragraph when trained and tested on each chain separately. Results are across the entire $F_v$ on model structures using our Expanded dataset.

### 7 Qualitative study - $F_{ab}$ fragment in complex with CD9 large extracellular loop

Figure S4 shows Paragraph's predictions for the PDB structure 6RLO compared to the ground truth. 6RLO is the structure of the CD9 large extracellular loop protein in complex with a neutralising antibody. 6RLO was chosen for this study to examine whether Paragraph is able to accurately predict the paratope when the antigen is bound away from the centre of the antibody. The figure shows how Paragraph successfully predicts the highest probabilities for the residues facing the antigen.

This example also highlights a limitation of our approach of predicting only residues within the CDR±2 and assigning zero probability to all other residues. The top two structures in Figure S4 show that Paragraph 'predicts' zero for two residues outside the CDR±2 that do belong to the paratope in this instance.

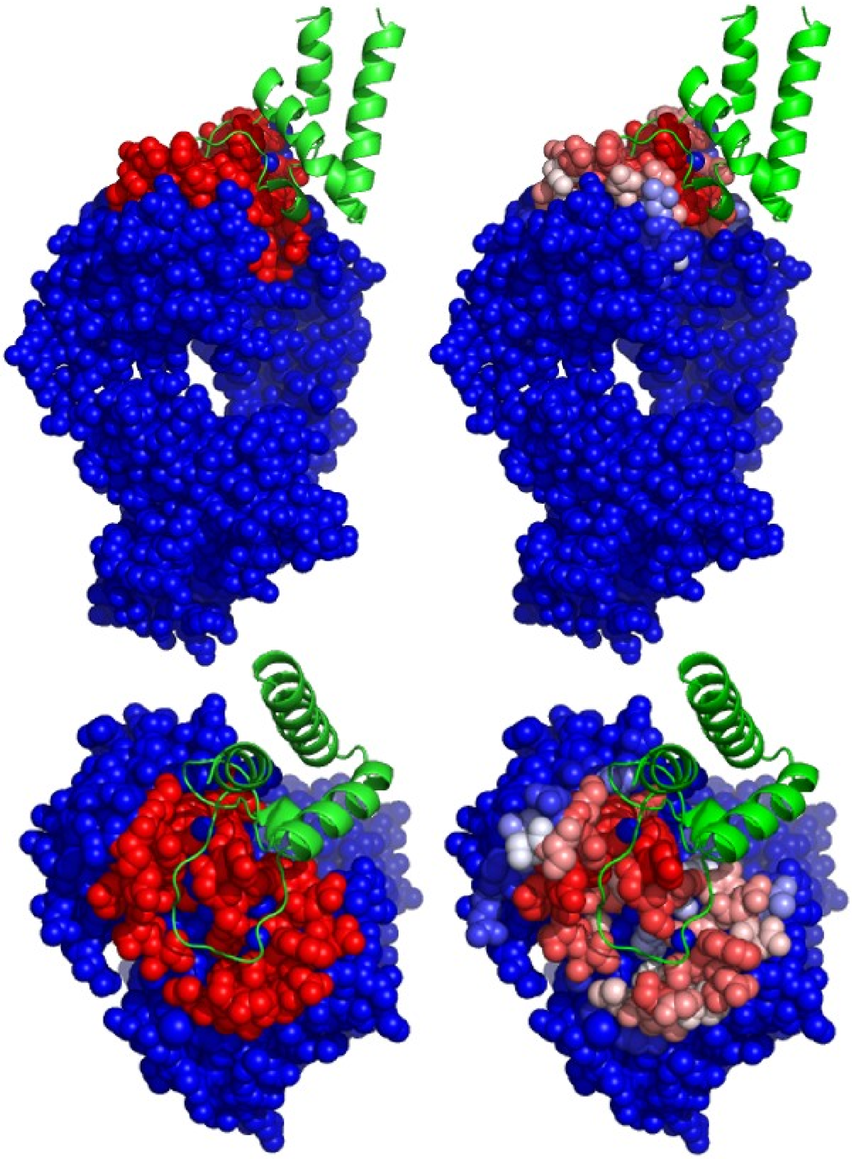

Figure S4: Side and top views of the $F_{ab}$ region of the CD9 large extracellular loop binding antibody, 6RLO. The left two figures show the ground truth paratope residues in red (within 4.5Å of an antigen heavy atom). Non-paratope residues are shown in blue. The antigen is shown as a cartoon representation in green. The right two figures show Paragraph's predicted paratope using a red-white-blue spectrum across the scores from Paragraph. Paragraph was trained on our Expanded dataset.

### 8 Paragraph performance over individual CDR loops and framework regions

A breakdown of Paragraph's performance when trained and tested on paired data in our Expanded dataset is provided below (Table S5). All results are provided on ABlooper[9] model structures of our test set and are evaluated over the entire $F_v$ region. In agreement with Parapred[6], thresholds for calculating the F-score and Matthews correlation coefficient (MCC) are obtained by maximising Youden's index[10].

Paragraph's performance over the framework region is lower than the CDR loops due to the larger class imbalance (very few framework residues belong to the paratope) and the fact that Paragraph is restricted to predicting the CDR±2 region only.

| Chain | PR AUC | ROC AUC | F-score | MCC |
|---|---|---|---|---|
| CDR-L1 | 0.762 | 0.857 | 0.678 | 0.499 |
| CDR-L2 | 0.675 | 0.876 | 0.654 | 0.559 |
| CDR-L3 | 0.770 | **0.884** | 0.747 | **0.598** |
| CDR-H1 | 0.735 | 0.856 | 0.678 | 0.515 |
| CDR-H2 | 0.789 | 0.854 | 0.727 | 0.498 |
| CDR-H3 | **0.796** | 0.866 | **0.762** | 0.571 |
| Framework | 0.429 | 0.768 | 0.505 | 0.500 |

Table S5: Performance of Paragraph broken down over individual CDR loops and the framework region. Results are across the entire $F_v$ on model structures using our Expanded dataset.

[3] James Dunbar, Konrad Krawczyk, Jinwoo Leem, Terry Baker, Angelika Fuchs, Guy Georges, Jiye Shi, and Charlotte M Deane. Sabdab: the structural antibody database. doi: 10.1093/nar/gkt1043. URL `www.opig.stats.ox.ac.uk/webapps/newsabdab/sabdab/`.

[4] Limin Fu, Beifang Niu, Zhengwei Zhu, Sitao Wu, and Weizhong Li. Cd-hit: accelerated for clustering the next-generation sequencing data. *Bioinformatics (Oxford, England)*, 28: 3150–3152, 12 2012. ISSN 1367-4811. doi: 10.1093/BIOINFORMATICS/BTS565. URL `https://pubmed.ncbi.nlm.nih.gov/23060610/`.

[5] Jinwoo Leem, James Dunbar, Guy Georges, Jiye Shi, and Charlotte M Deane. mabs abodybuilder: Automated antibody structure prediction with data-driven accuracy estimation abodybuilder: Automated antibody structure prediction with data-driven accuracy estimation. 2016. doi: 10.1080/19420862.2016.1205773. URL `https://www.tandfonline.com/action/journalInformation?journalCode=kmab20`.

[6] Edgar Liberis, Petar Veli Ckovi C, Pietro Sormanni, Michele Vendruscolo, and Pietro Liò. Parapred: antibody paratope prediction using convolutional and recurrent neural networks. doi: 10.1093/bioinformatics/bty305. URL `http://www-mvsoftware.ch.cam.ac.uk/andfordownloadathttps://github.com/eliberis/parapred`.

[7] Phil Wang LucidRains. Egnn - pytorch, 2021. URL `www.github.com/lucidrains/egnn-pytorch`.

[8] Victor Garcia Satorras, Emiel Hoogeboom, and Max Welling. E(n) equivariant graph neural networks, 2021.

[9] Brennan Abanades, Guy Georges, Alexander Bujotzek, and Charlotte M Deane. Ablooper: Fast accurate antibody cdr loop structure prediction with accuracy estimation. 2021. doi: 10.1101/2021.07.26.453747. URL `https://doi.org/10.1101/2021.07.26.453747`.

[10] W. J. Youden. Index for rating diagnostic tests. *Cancer*, 3(1):32–35, 1950. doi: https://doi.org/10.1002/1097-0142(1950)3:1⟨32::AID-CNCR2820030106⟩3.0.CO;2-3.

[11] Marie-Paule Lefranc, Vé ronique Giudicelli, Chantal Ginestoux, Joumana Jabado-Michaloud, Gé raldine Folch, Fatena Bellahcene, Yan Wu, and Elodie Gemrot. Imgt, the international immunogenetics information system. *Nucleic Acids Research*, 37, 2009. doi: 10.1093/nar/gkn838. URL `https://academic.oup.com/nar/article/37/suppl_1/D1006/1014137`.
